# Reducing oxidative stress improves ex vivo polymer-based human haematopoietic stem and progenitor cell culture and gene editing

**DOI:** 10.1101/2024.09.17.613552

**Authors:** Yavor Bozhilov, Elizabeth Brown, Ian Hsu, Indranil Singh, Alejo Rodriguez-Fraticelli, Anindita Roy, Satoshi Yamazaki, Adam C. Wilkinson

## Abstract

Self-renewing multipotent haematopoietic stem cells (HSCs) have the unique capacity to stably regenerate the entire blood and immune systems following transplantation. HSCs are used clinically to reconstitute a healthy blood system in patients suffering from a range of haematological diseases. However, HSCs are very rare and have been challenging to grow ex vivo, which has hampered efforts to collect large numbers of HSCs for both basic research and clinical therapies. Polymer-based culture conditions have recently been developed to support expansion of mouse and human haematopoietic stem and progenitor cells (HSPCs). While mouse HSPCs expanded rapidly in polymer-based cultures, growth speeds for human HSPCs in polymer-based cultures was limited to ∼70-fold over 4-weeks. Here we have found that reducing oxidative stress improves human HSPC growth in these conditions. We describe an optimised culture condition that improves growth to 250-1400-fold over 4-weeks through reducing oxygen tension and the addition of antioxidants. These conditions also enable efficient gene editing in these polymer-based cultures. We envision these improved culture conditions will support a range of research into human HSPC biology and provide a platform for clinical-scale HSPC expansion and gene editing.

## Introduction

Self-renewing multipotent haematopoietic stem cells (HSCs) are a very rare cell type with the unique capacity to stably regenerate the entire blood and immune system^1, 2^. HSC transplantation has been used clinically for over 60 years in the treatment of haematological malignancies^3^, and has more recently been combined with gene therapies to treat a range of non-malignant disorders^4^. HSC dose can limit clinical use and patient safety, particularly for allogeneic HSC transplantation that uses umbilical cord blood (UCB) as a donor HSC source^5^, and HSC gene therapies where toxicities associated with ex vivo gene editing can result in delayed engraftment^6^.

While HSCs survive and retain their functional potential throughout life within their endogenous in vivo bone marrow microenvironment, HSCs easily lose their function when placed in ex vivo culture^7^. This has hampered efforts to improve the safety and availability of HSC cell and gene therapies. Lack of culture conditions to stably grow functional HSCs long-term ex vivo has also limited our ability to study this biologically and clinically important stem cell population. Over the years, several culture systems have been developed^7^, with some tested clinically for expanding UCB haematopoietic stem and progenitor cells (HSPCs)^8, 9^. However, these conditions generally only supported short-term maintenance and expansion.

We previously developed a polymer-based culture condition that could support long-term and large-scale expansion of transplantable mouse HSPCs^10, 11^. Adaption of these conditions for human UCB HSPCs led to a new chemically-defined “3a” media that supported transplantable human HSPCs for at least one month^12^. Stable growth conditions for human HSPCs have abundant opportunities to support improve clinical cell and gene therapies, and to study human HSPC biology and haematopoiesis. However, limiting the utility of the current system, human UCB-derived HSPCs grew slowly in these conditions (∼70-fold expansion over 30-days). Additionally, while there is substantial interest in the scientific and clinical applications of CRISPR/Cas9-based HSC gene editing^4^, CRISPR/Cas9 has not yet been applied to these polymer-based human HSPC cultures.

Here, we sought to optimise polymer-based 3a media conditions to enhance human UCB HSPC growth ex vivo through modulating oxidative stress. Our optimised conditions boost cell expansion from ∼70-fold to ∼400-fold, while retaining equivalent engraftment potential and clonality, and significantly improving HSPC survival during ex vivo gene editing.

## Materials and Methods

### Human HSPCs

Primary human cells were used with informed consent and ethical approval (OxTREC Ref. 573-23 and REC Ref. 21/LO/0195). Fresh human UCB CD34^+^ cells (provided by NHSBT) were isolated using a Ficoll gradient and the Miltenyi CD34-enrichment microbead kit. Alternatively, cryopreserved CD34^+^ cells were purchased from StemExpress (Folsom). CD34^+^ cells were cryopreserved in liquid nitrogen until use.

### Cell cultures

CD34^+^ HSPCs were cultured in IMDM media (Gibco cat# 12440-053) supplemented with penicillin-streptomycin-glutamine (1X; Gibco cat# 10378-016), insulin-transferrin-selemium-ethanolamide (1X; Gibco cat# 51500-056), Soluplus (0.1% w/v; BASF cat# 50539897), 740 Y-P (5 μM; MedChemExpress cat# HY-P0175), butzyamide (100 nM; a kind gift from Celaid), UM729 (1.5 μM; StemCell Technologies cat# 72332), recombinant human FLT3L (10 ng/ml; Peprotech cat# AF-300-19), and/or monothioglyercol (100 μM; Sigma cat# M1753). Where indicated, cells were cultured with beta-mercaptoethanol, N-acetyl cysteine, ascorbic acid, ascorbic acid 2-phosphate, Trolox, LCS1, or DPI (Sigma). 10,000 CD34^+^ cells were seeded per well in 96-well CellBIND plates (Corning cat# 3300) and incubated at 37°C with 5% CO_2_ and the indicated O_2_ level. Compete media changes were performed every 2-3 days. For K562 and OCI-AML3 leukaemia cell lines, cells were cultured in IMDM supplemented with 10% fetal bovine serum (Gibco) and penicillin-streptomycin-glutamine.

### Flow cytometric analysis

At indicated timepoints, cells were collected and stained with anti-human CD34-APC/Cy7, CD90-FITC, CD201-PE, CD45RA-PacificBlue, CD41-APC (BioLegend), and propidium iodide (PI). Flow cytometry was performed using a BD LSRFortessa. For the intracellular ROS assay, cultured cells were stained with dihydroethidium prior to flow cytometric analysis. For bromodeoxyuridine (BrdU) incorporation assays, BrdU was added to cell cultures 30 minutes prior to collection and fixation using the BD cytofix/cytoperm kit and staining with an anti-BrdU-APC (Biolegend) antibody. For the intracellular cell cycle analysis, cultured cells were fixed using the BD cytofix/cytoperm kit and stained with anti-Ki67-PE (BD) and anti-phospho-histone H3 (Ser10)-FITC (Cell Signalling) antibodies and Hoescht 33342 (ThermoFisher Scientific).

### Colony forming unit (CFU) assays

CFU assays were performed using Methocult (H4435; StemCell Technologies). 500 cells were plated in 1 ml Methocult according to the manufacturer’s instructions and incubated at 37°C with 5% CO_2_ for 12-14 days before CFUs were scored.

### Xenotransplantation assays

All animal experiments were performed in accordance with institutional guidelines and were approved by the UK Home Office and the University of Oxford AWERB committee. Immunodeficient NOD.Cg-Prkdc^scid^ Il2rg^tm1Wjl^/SzJ (NSG) mice were bred at the University of Oxford. Human HSPCs were transplanted into 8-16-week-old recipient mice (following 2 Gy sub-lethal irradiation) and human chimerism determined by analysis with anti-human CD45-PacificBlue, anti-human HLA-ABC-FITC, and anti-mouse CD45.1-PE/Cy7.

### Barcoding assay

We used the pLARRYv2 lentiviral library^13^ to transduce 200,000 fresh CD34^+^ HSPCs at an MOI of 0.1 to barcode ∼20,000 HSPCs. HSPCs were then split in the conditions indicated (∼10,000 barcoded cells per condition). The cells were in culture for 28-days before gDNA extraction, barcode library preparation, and NGS sequencing. Sequencing reads were initially screened for quality using fastQC, then the reads were filtered and trimmed down to the barcode region using trimmomatic, trim_galore and cutadapt. To account for sequencing error sequences were collapsed with Hamming distance of 4 then unique sequences were extracted and counted.

### Transcriptomic analysis

Indicated cell culture populations were isolated by FACS and RNA extracted. RNA library preparation and sequencing was performed by Novogene. Sequencing reads were initially screened for quality using fastQC. Subsequently, paired sequencing reads were aligned to the human genome using STAR^14^, and raw gene counts were generated using subread featureCounts^15^. The gene counts were passed into DESeq2 workflow^16^ for normalisation, and PCA was generated using the prcomp function. Contrasts were created to perform Wald testing on shrunken log2 fold changes (type = ashr), the outputs of which were used for volcano plots. Significant differentially expressed genes (padj <0.01, log2FoldChange > |1|) were used for gene set enrichment analysis.

### Gene editing

Human HSPCs were cultured for 4-7 days and then collected for gene edited. HSPCs were electroporated with pre-complexed Cas9/sgRNA ribonucleoprotein (RNP) complexes using the Lonza Nucleofector 4D system using programme EO-100 and primary P3 buffer. Cas9/sgRNA RNPs were complexed by mixing recombinant Cas9 (IDT) with synthetic sgRNAs (IDT) for 15 minutes at 20°C. HSPCs were immediately returned to wells containing fresh media and analysed at 3-days post-electroporation.

## Results

Given our previous success with improving mouse polymer-based HSC cultures using 5% O_2_, we wanted to evaluate how reducing oxygen tension influenced human HSPC growth in similar conditions. However, we first needed to establish the 3a culture media with commercially-available reagents. We therefore evaluated UM729 as an alternative to UM171^17^. While total cell expansion was slightly reduced with UM729 after 14 days (**Figure S1A**), there was an equivalent number of CD34^+^CD90^+^CD201^+^CD45RA^-^CD41^-^ cells (**Figure S1B**), a reported ex vivo immunophenotypic (p)HSC population^18^. By contrast, the number of CD41^+^ (megakaryocyte lineage) cells were significantly reduced by UM729 (**Figure S1C**). We could further inhibit growth of CD41^+^ cells by the addition of 10 ng/ml FLT3L (**Figure S1D**), consistent with a recent report^19^. Using this updated media cocktail, we evaluated human CD34^+^ UCB HSPC growth at various oxygen levels (**Figure 1A**). Cell growth and frequency of pHSCs were similar between 20% and 5% O_2_ (**Figure 1B-C**). However, when cells were grown at 10% O_2_, cell growth approximately doubled whilst maintaining similar pHSC frequencies. These results suggested that reducing oxygen tension could improve human HSPC growth in polymer-based cultures, but that a certain level of oxygen was needed for cell proliferation.

**Figure 1.**
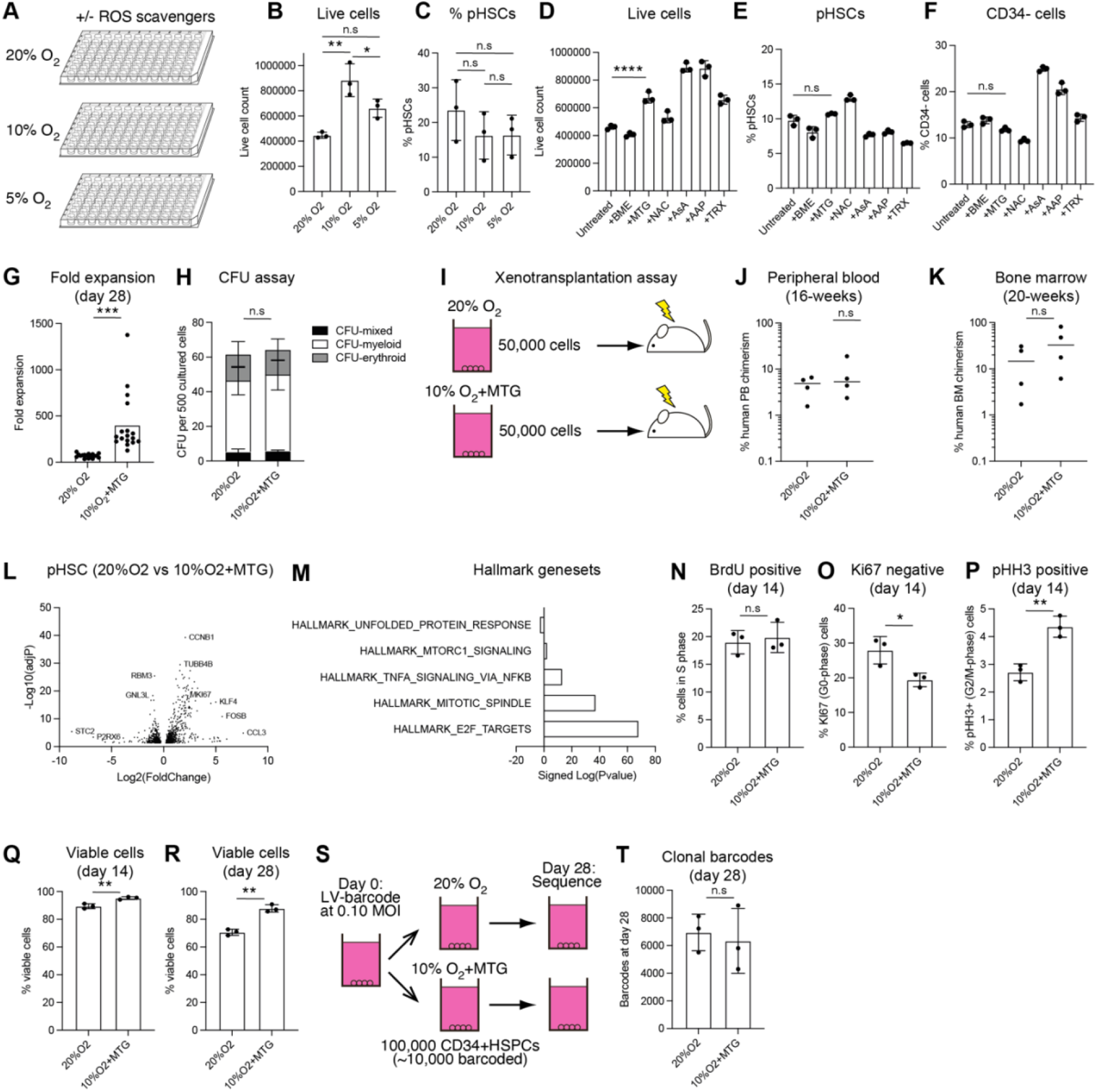
Improved polymer-based human HSPC survival and expansion through reducing oxidative stress. a) Schematic of titrating O_2_tension and ROS scavengers for human UCB CD34^+^ HSPC cultures. b) Average number of live cells derived from 10,000 UCB CD34^+^ HSPCs cultured as indicated in (a) for 14-days. Data from 3 biological replicates. Statistical analysis by one-way ANOVA. c) Average frequency of immunophenotypic CD34^+^CD90^+^CD201^+^CD45RA^-^CD41^-^ (p)HSCs derived from 10,000 UCB CD34^+^ HSPCs cultured as indicated in (a) for 14-days. Data from 3 technical replicates. Statistical analysis by one-way ANOVA. d) Average number of live cells derived from 10,000 UCB CD34^+^ HSPCs cultured with indicated ROS scavengers for 14 days at 10% O_2_; beta-mercaptoethanol (BME), monothioglycerol (MTG), N-acetyl cysteine, ascorbic acid (AsA), ascorbic acid 2-phosphate (AAP), and Trolox (TRX). Data from 3 technical replicates. Statistical analysis by one-way ANOVA. e) Average frequency of pHSCs derived from 10,000 UCB CD34^+^ HSPCs cultured with indicated ROS scavengers for 14 days at 10% O_2_. Data from 3 technical replicates. Statistical analysis by one-way ANOVA. f) Average frequency of immunophenotypic CD34^-^ cells derived from 10,000 UCB CD34^+^ HSPCs cultured with indicated ROS scavengers for 14 days at 10% O_2_. Data from 3 technical replicates. Statistical analysis by one-way ANOVA. g) Average fold change in live cell numbers after a 28-day CD34^+^ HSPC culture using 20%O_2_ or 10%O_2_+MTG conditions. Data from 17 biological replicates. Statistical analysis by paired T-test. h) Average number of colony-forming units (CFU) generated from 500 14-day cultured cells from 20%O_2_ or 10%O_2_+MTG conditions. Data from 3 biological replicates. Statistical analysis by paired T-test. i) Schematic for xenotransplantation assay using 50,000 14-day cultured cells. j) 16-week human CD45^+^ chimerism in the peripheral blood of recipient mice. Data from 4 recipient mice per cohort. Statistical analysis by unpaired T-test. k) 20-week human CD45^+^ chimerism in the bone marrow of recipient mice. Data from 4 recipient mice per cohort. Statistical analysis by unpaired T-test. l) Differentially expressed genes between pHSCs from 20%O_2_ and 10%O2+MTG cultures at day 14 displayed as Log2(fold change) vs -Log10(adjusted P value). RNA sequencing performed on 3 biological replicates. m) Signed Log(p-values) for gene set enrichment within Hallmark gene sets. n) Frequency of BrdU positive cells in 14-day HSPC cultures, following a 30-minute labelling with BrdU. Data from 3 biological replicates. Statistical analysis by paired T-test. o) Frequency of KI67 negative cells in 14-day HSPC cultures. Data from 3 biological replicates. Statistical analysis by paired T-test. p) Frequency of phospho-histone H3 (pHH3-Ser10) positive cells in 14-day HSPC cultures. Data from 3 biological replicates. Statistical analysis by paired T-test. q) Frequency of viable (PI-negative Annexin V-negative) cells in 14-day cultures. Data from 3 biological replicates. Statistical analysis by paired T-test. r) Frequency of viable (PI-negative Annexin V-negative) cells in 28-day cultures. Data from 3 biological replicates. Statistical analysis by paired T-test. s) Schematic for clonal barcoding analysis of HSPC cultures. t) Number of barcodes detected in 28-day cultures, initiated with 10,000 clonally-barcoded HSPCs. Data from 3 biological replicates. Statistical analysis by paired T-test. For all statistical analysis, * denotes p<0.05; ** denotes p<0.005; *** denotes p<0.0005; **** denotes p<0.0001.

We next hypothesised that addition of antioxidants could be used to reduce oxygen-related cellular stresses and thereby further boost cell growth. We screened six antioxidants in our CD34^+^ UCB HSPC cultures; beta-mercaptoethanol (BME), monothioglycerol (MTG), N-acetyl cysteine (NAC), ascorbic acid (AsA), ascorbic acid 2-phosphate (AAP), and Trolox (TRX). Of these antioxidants, MTG, AsA, AAP and TRX further boosted total cell growth within the 10% O_2_ cultures (**Figure 1D**). However, only MTG cultures retained similar frequencies of pHSCs (**Figure 1E**). The reductions in pHSC frequency in Asa, AAP and TRX cultures were likely due to greater differentiation, as higher frequencies of CD34^-^ cells were observed within the cultures (**Figure 1F**). We therefore selected MTG as the optimal additive for 10% O_2_ cultures to maximise HSPC expansion. Compared to the 20%O_2_ cultures, our optimised 10%O_2_+MTG cultures generated an average of ∼400-fold cell growth over 28-days (**Figure 1G**), over 5-times more than achieved in 20%O_2_ cultures. It is worth noting that MTG also improved HSPC expansion in 20%O_2_ cultures, although not to the extent of the 10%O_2_+MTG cultures (**Figure S1E**).

To initially confirm that we retained equivalent HSPCs within these optimised 10%O_2_+MTG cultures, we performed colony forming unit (CFU) assays (**Figure 1H**). 10%O_2_+MTG cultures generated a range of myeloid, erythroid, and mixed colonies, suggesting a similar functional composition of HSPCs within the cultures. Next, to confirm the engraftment potential of these cultured cells, we performed xenotransplantation assays (**Figure 1I**). We transplanted 50,000 cells derived from either 20%O_2_ or 10%O_2_+MTG cultures into sub-lethally irradiated immunodeficient NSG mice. Similar levels of human chimerism were seen both in the peripheral blood at 16-weeks and in the bone marrow at 20-weeks (**Figure 1K**). Equivalent results were also achieved with two other UCB samples (**Figure S1F-G**). We therefore concluded that use of 10%O_2_+MTG enhanced HSPC growth by ∼5-fold within these cultures without substantially altering the type of cells generated in these conditions.

To evaluate whether our 10%O_2_+MTG cultures modified molecular features of the HSPCs in these cultures, we performed RNA sequencing on CD201^+^CD90^+^CD34^+^CD41^-^ pHSCs and CD201^-^ haematopoietic progenitor cells (pHPCs) from three different donors. Principal component analysis confirmed that pHSCs and pHPCs separated from each other, although also highlighted donor-specific effects (**Figure S1H**). More differentially expressed genes (DEGs) were observed between pHSCs and pHPCs (2380 genes for 10%O_2_+MTG cultures; 1793 genes for 20%O_2_ cultures) than between pHSCs from the different conditions (703 genes) (**Figures 1L, S1I-J**). pHSC-specific genes included known HSC markers^11^ including *HLF, MECOM, FGD5*, and *MLLT3*. Focusing on the 259 statistically significant DEGs (padj < 0.01) with over 2-fold changes (199 genes upregulated, 60 genes downregulated), gene set enrichment analysis identified genes associated with cell division, TNFA signalling, and MTOR signalling as upregulated in 10%O_2_+MTG cultures, while genes associated with the unfolded protein response were upregulated in 20%O_2_ cultures (**Figure 1M**).

To better understand how 10%O_2_+MTG influenced cell cycling of HSPCs ex vivo, we quantified cell cycle by flow cytometry. We were unable to identify differences in the frequency of S-phase cells between 20%O_2_ and 10%O_2_+MTG cultured cells using BrdU labelling (**Figure 1L**). However, we could identify a modest decrease in Ki67 negative (putative G0 cells) and a modest increase in phospho-histone H3 positive (putative G2/M cells) in 10%O_2_+MTG cultures (**Figure 1O-P**). These differences likely contributed to the gene expression differences observed in the transcriptional analysis but seem less likely to drive the differences in fold expansion. As enhanced cell expansion could also be driven by improved cell survival, we next quantified apoptosis using Annexin-V. We could identify significantly higher rates of cell death at 20%O_2_, particularly by day 28 (**Figure 1Q-R**). These results suggest that 10%O_2_+MTG may therefore enhance overall cell expansion by promoting cell cycle entry and cell survival.

The improved HSPC expansion in the 10%O_2_+MTG cultures could be due to altered clonal outgrowth. To better understand whether cell death within the culture is random, or represented a loss of clonality, we performed clonal barcoding analysis using the pLARRY^20, 21^ lentiviral barcode system (**Figure 1S**). After transducing ∼10,000 fresh CD34^+^ HSPCs per condition, we cultured these HSPCs for 28-days at 20%O_2_ or 10%O_2_+MTG, and then quantified barcodes still present in each culture condition. We observed very similar clonal outgrowth in 20%O_2_ and 10%O_2_+MTG cultures, recovering ∼6,000 barcodes from both conditions (**Figure 1T**). We therefore conclude that these long-term cultures do not induce a high degree of selective pressure or oligoclonal outgrowth.

Together, the results above suggested that the improved HSPC expansion in polymer-based cultures achieved using 10%O_2_+MTG was due to reducing oxidative stress. To better investigate the consequence of 10%O_2_ and MTG on intracellular ROS accumulation in these human HSPC cultures, we quantified intracellular ROS levels (**Figure 2A**). Surprisingly, we observed modest increases in ROS levels in the 10%O_2_+MTG cultured HSPCs, suggesting that rather than quenching ROS, 10%O_2_+MTG may act as a buffer to allow human HSPCs to adapt and survive higher levels of intracellular ROS. However, inhibition of superoxide dismutase (SOD1) using LCS1 was highly toxic to human HSPCs in these cultures (**Figure 2B-C**), implying that these HSPCs cannot accommodate further increases in ROS. Of note, the same dose of LCS1 was well tolerated by human leukaemia cell lines (OCI-AML1 and K562) (**Figure 2C**), suggesting that the balance between ROS and antioxidants is much more tightly regulated in HSPCs.

**Figure 2.**
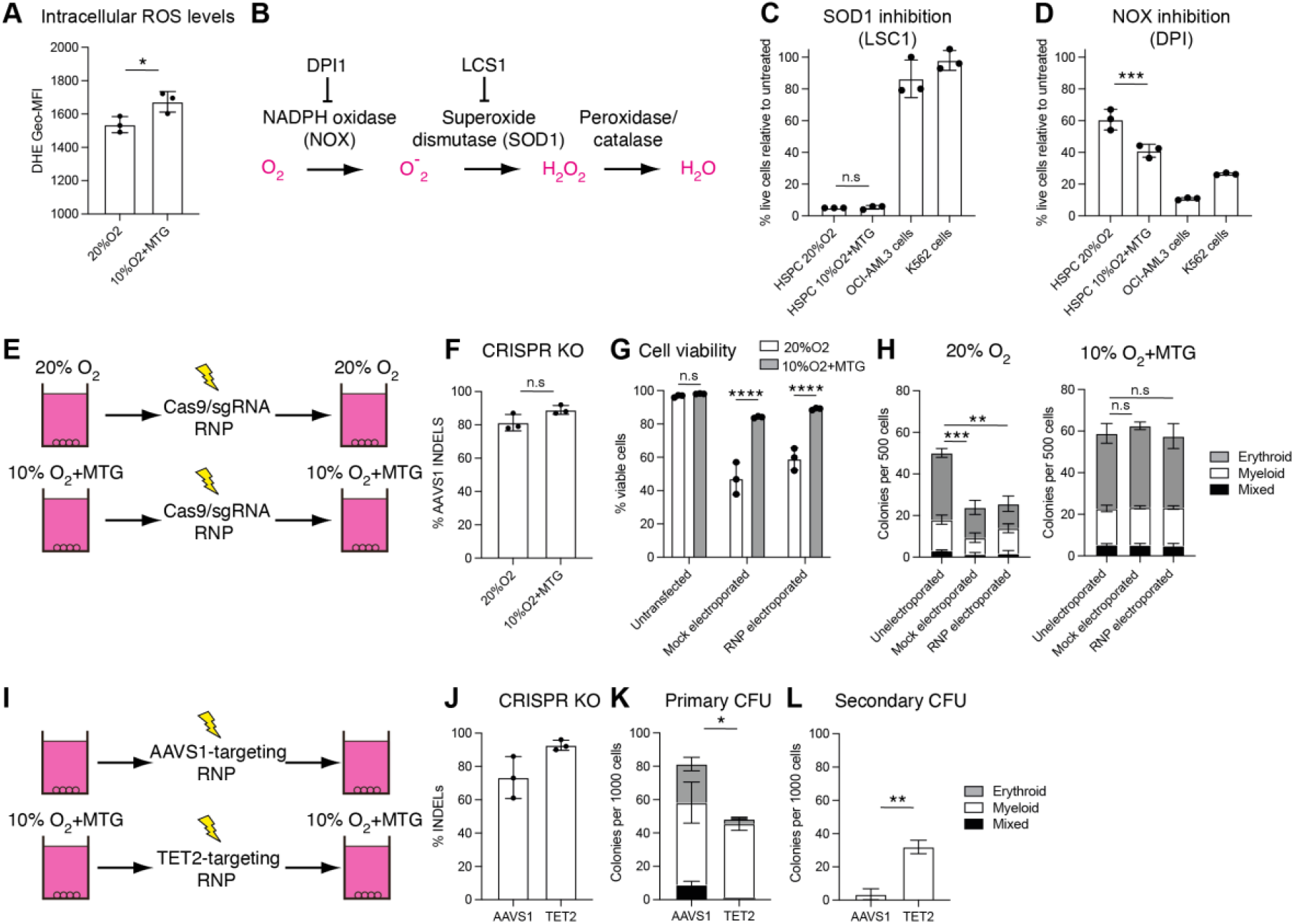
Reduced oxidative stress in ex vivo cultures limits HSPC loss during gene editing. a) Dihydroethidium (DHE) geometric mean fluorescence intensity (MFI) as measure of intracellular ROS levels for human HSPCs cultured in the indicated conditions. Data from 3 biological replicates. Statistical analysis by paired T-test. b) Schematic of intracellular ROS generation via NADPH oxidase (NOX) and breakdown by superoxide dismutase (SOD1), with inhibitors used in this study indicated: LSC1 and diphenyleneiodonium chloride (DPI). c) Relative growth of human HSPCs or OCI-AML3 and K562 cell lines cultured with 1 μM LCS1, normalised to DMSO control cultures. Data from 3 biological replicates. Statistical analysis by one-way ANOVA. d) Relative growth of human HSPCs or OCI-AML3 and K562 cell lines cultured with 250 nM DPI, normalised to DMSO control cultures. Data from 3 biological replicates. Statistical analysis by one-way ANOVA. e) Schematic of CRISPR/Cas9-based gene editing of the *AAVS1* safe harbour loci in 20%O_2_ and 10%O_2_+MTG cultured HSPCs. f) Frequency of *AAVS1* indels at 3-days post-electroporation of Cas9/sgRNA as indicated in (e). Data from 3 biological replicates. g) Frequency of viable (PI-negative Annexin V-negative) cells at 3-days post-electroporation of Cas9/sgRNA as indicated in (e). Data from 3 biological replicates. Statistical analysis by one-way ANOVA. h) Number of CFUs generated from 500 live cells isolated 3-days post-electroporation of Cas9/sgRNA as indicated in (e). Data from 3 biological replicates. Statistical analysis by one-way ANOVA. i) Schematic of CRISPR/Cas9-based genetic mutation of *TET2* or *AAVS1* in 10%O_2_+MTG cultured HSPCs. j) Frequency of indels at 3-days post-electroporation of Cas9/sgRNA, as indicated in (i). Data from 3 biological replicates. k) Number of primary CFUs generated from 1000 live cells isolated 3-days post-electroporation with TET2 or *AAVS1*-targeting Cas9/sRNA, as indicated in (i). Data from 3 biological replicates. Statistical analysis by paired T-test. l) Number of secondary CFUs generated from replating 1000 cells from the primary CFU detailed in (k). Data from 3 biological replicates. Statistical analysis by paired T-test. For all statistical analysis, * denotes p<0.05; ** denotes p<0.005; *** denotes p<0.0005; **** denotes p<0.0001.

We also investigated the consequences of reducing intracellular ROS via inhibition of NADPH oxidase (NOX) using the small molecule diphenyleneiodonium chloride (DPI) (**Figure 2B**). Interestingly, MTG-cultured cells displayed enhanced sensitivity to DPI (**Figure 2D**). These results suggest that the higher ROS levels in the 10%O_2_+MTG cultures may be important for HSC proliferation. Fast-growing human leukaemic (OCI-AML1 and K562) cell lines were also highly sensitive to DPI (**Figure 2D**). Together, these results suggest that while human HSPCs are sensitive to high intracellular ROS, ROS is also important for cell proliferation. Future studies will be needed to understand the molecular rheostat acting in human HSPCs to finetune ROS levels.

Next, we wanted to investigate the application of these cultures to perform electroporation-based HSPC gene editing. Electroporation has been shown to induce ROS^22^, leading us to hypothesise that 10%O_2_+MTG culture may improve survival in the context of electroporation-based CRISPR/Cas9 gene editing. We therefore evaluated CRISPR/Cas9-based gene editing of HSPCs from 20%O_2_ or 10%O_2_+MTG cultures (**Figure 2E**). In both conditions, we could achieve high rates of editing by electroporation of pre-complexed recombinant Cas9 and synthetic sgRNA targeting the *AAVS1* safe harbour loci^23^ (**Figure 2F**). However, survival was superior in the 10%O_2_+MTG cultures (**Figure 2G**). Additionally, when clonal potential was evaluated by CFU assays, the HSPCs that had survived gene editing from the 20%O_2_ culture cells also displayed loss of CFU potential (**Figure 2H**). Taken together, these results highlight the significant benefit of using 10%O_2_+MTG conditions for gene editing human HSPCs in these polymer-based cultures.

Finally, as a proof-of-concept for the use of this system to study HSPC biology, we mutated *TET2* using a previously reported sgRNA^24^ (**Figure 2I**). We again achieved high rates of editing (**Figure 2J**). Additionally, consistent with the reported consequence of loss of *TET2* in HSPCs^24, 25^, serial replating CFU assays confirmed that *TET2*-mutant HSPCs displayed a loss of erythroid potential (**Figure 2K**) while exhibiting enhanced myeloid replating capacity (**Figure 2L**). Together, these results validate our 10%O_2_+MTG culture conditions as highly supportive of human HSPC expansion and gene editing.

## Summary

In summary, we have developed an optimised ex vivo polymer-based culture condition for the long-term growth and gene editing of human UCB HSPCs. We achieved these improvements using 10% O_2_ and the addition of the antioxidant MTG. Our proof-of-concept gene editing results also highlight the potential utility of this optimised culture system to study the biology of human HSPCs and model the consequences of genetic mutations within a tractable ex vivo system. We envision that these approaches will have wide applications for the haematology and stem cell communities.

## Acknowledgements

We thank the WIMM Flow Cytometry Core for flow cytometry support. This research was supported by grants from the Kay Kendall Leukaemia Fund, the MRC, NIHR Oxford-Birmingham Blood and Transplant Research Unit in Advanced Cellular Therapeutics, NIHR Oxford Biomedical Research Centre, the John Fell Fund, an Oxford Goodger and Schorstein Scholarship, and the Oxford Cancer Centre.

## Authorship statement

Y.B., I.H., E.B., I.S., A.R.F, A.R, and S.Y performed experiments, provided reagents, and edited the manuscript. Y.B. and A.C.W. analysed the data and wrote the manuscript.

---

**Supplementary Figure 1.**
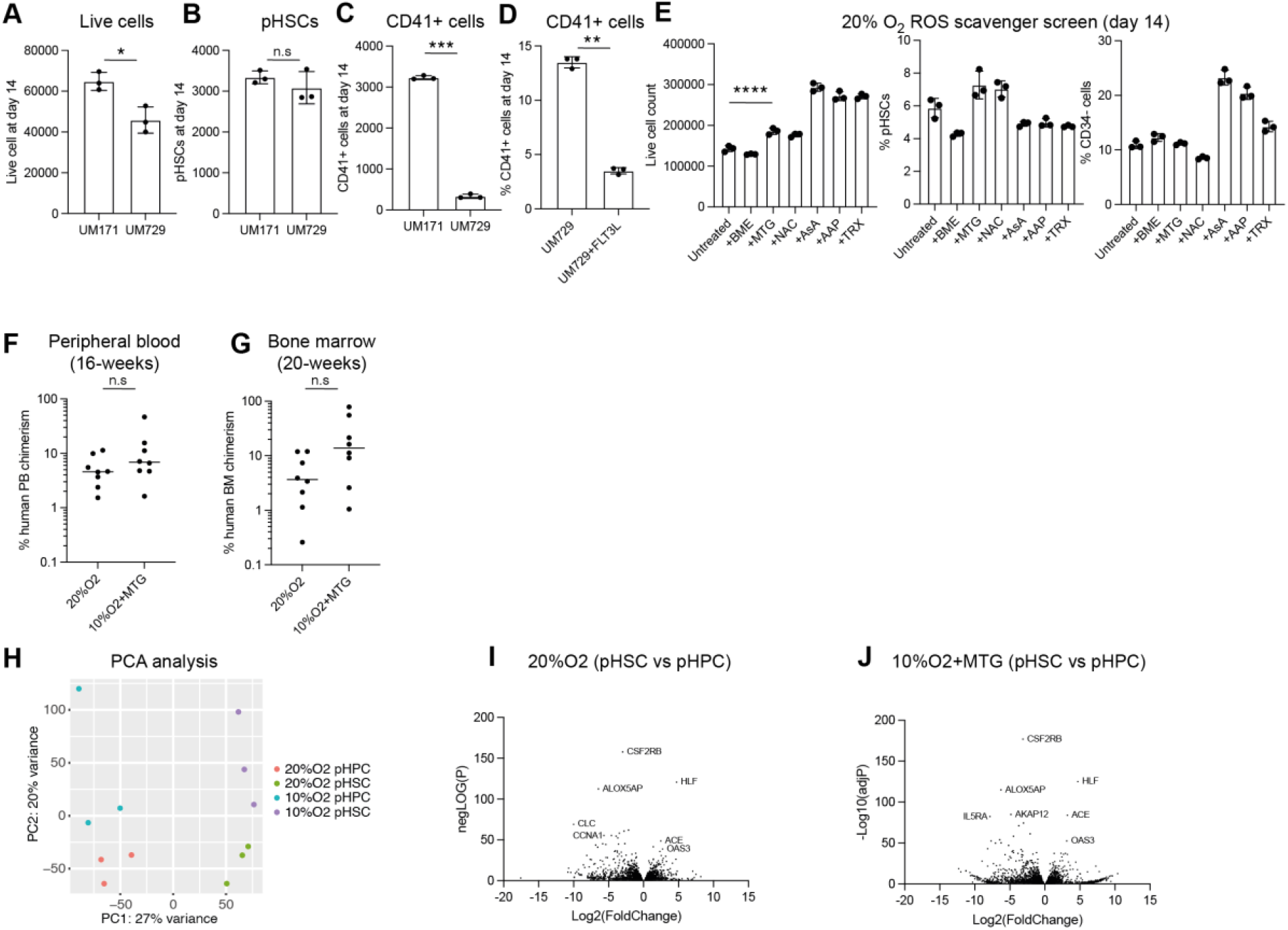
Related to Figure 1. a) Number of cells generated from 10,000 CD34^+^ HSPCs in 14-day cultures supplemented with butyzamide, 740Y-P and the addition of UM171 or UM729. Data from 3 biological replicates. Statistical analysis by paired T-test. b) Number of immunophenotypic CD34^+^CD90^+^CD201^+^CD45RA^-^CD41^-^ (p)HSCs derived from 10,000 UCB CD34^+^ HSPCs cultured as indicated in (a). Data from 3 biological replicates. Statistical analysis by paired T-test. c) Number of CD41^+^ megakaryocyte-lineage cells generated from 10,000 UCB CD34^+^ HSPCs cultured as indicated in (a). Data from 3 biological replicates. Statistical analysis by paired T-test. d) Frequency of CD41^+^ megakaryocyte-lineage cells generated in 14-day cultures supplemented with butyzamide, 740Y-P, UM729, and with or without addition of FLT3L. Data from 3 biological replicates. Statistical analysis by paired T-test. e) ROS scavenger screen (as described in **Figure 1d**) performed at 20% O_2_, with number of live cells (left panel), % pHSCs (middle panel) and % CD34-negative cells (right panel) at day 14 displayed. Data from 3 technical replicates. Statistical analysis by one-way ANOVA. f) 16-week human CD45^+^ chimerism in the peripheral blood of recipient mice from 100,000 day-14 cultured cells. Data from 8 recipient mice per cohort from two independent UCB samples. Statistical analysis by unpaired T-test. g) 20-week human CD45^+^ chimerism in the bone marrow of recipient mice from 100,000 day-14 cultured cells. Data from 8 recipient mice per cohort from two independent UCB samples. Statistical analysis by unpaired T-test. h) Principal component analysis of RNA-seq samples from pHSC and pHPC populations from 20%O_2_ and 10%O_2_+MTG cultures collected at day 14. i) Differentially expressed genes between pHSCs and pHPCs from 20%O_2_ cultures at day 14 displayed as Log2(fold change) vs -Log10(adjusted P value). RNA sequencing performed on 3 biological replicates. j) Differentially expressed genes between pHSCs and pHPCs from 10%O_2_+MTG cultures at day 14 displayed as Log2(fold change) vs -Log10(adjusted P value). RNA sequencing performed on 3 biological replicates. For all statistical analyses, * denotes p<0.05; ** denotes p<0.005; *** denotes p<0.0005; **** denotes p<0.0001.

## References

1. Eaves, C.J. Hematopoietic stem cells: concepts, definitions, and the new reality. Blood 125, 2605–2613 (2015).

2. Wilkinson, A.C., Igarashi, K.J. & Nakauchi, H. Haematopoietic stem cell self-renewal in vivo and ex vivo. Nat Rev Genet (2020).

3. Chabannon, C. et al. Hematopoietic stem cell transplantation in its 60s: A platform for cellular therapies. Sci Transl Med 10 (2018).

4. Ferrari, S. et al. Genetic engineering meets hematopoietic stem cell biology for next-generation gene therapy. Cell Stem Cell 30, 549–570 (2023).

5. Ballen, K.K., Gluckman, E. & Broxmeyer, H.E. Umbilical cord blood transplantation: the first 25 years and beyond. Blood 122, 491–498 (2013).

6. Frangoul, H. et al. CRISPR-Cas9 Gene Editing for Sickle Cell Disease and beta-Thalassemia. N Engl J Med 384, 252–260 (2021).

7. Meaker, G.A. & Wilkinson, A.C. Ex vivo hematopoietic stem cell expansion technologies: recent progress, applications, and open questions. Exp Hematol 130, 104136 (2023).

8. Cohen, S. et al. Hematopoietic stem cell transplantation using single UM171-expanded cord blood: a single-arm, phase 1–2 safety and feasibility study. Lancet Haematol 7, e134–e145 (2020).

9. Horwitz, M.E. et al. Phase I/II Study of Stem-Cell Transplantation Using a Single Cord Blood Unit Expanded Ex Vivo With Nicotinamide. J Clin Oncol 37, 367–374 (2019).

10. Wilkinson, A.C. et al. Long-term ex vivo haematopoietic-stem-cell expansion allows nonconditioned transplantation. Nature 571, 117–121 (2019).

11. Igarashi, K.J. et al. Physioxia improves the selectivity of hematopoietic stem cell expansion cultures. Blood Adv (2023).

12. Sakurai, M. et al. Chemically defined cytokine-free expansion of human haematopoietic stem cells. Nature 615, 127–133 (2023).

13. Singh, I., Fernandez Perez, D., Sánchez, P. & Rodriguez-Fraticelli, A.E. Pre-existing stem cell heterogeneity dictates clonal responses to acquisition of cancer driver mutations. bioRxiv, 2024.2005.2014.594084 (2024).

14. Dobin, A. et al. STAR: ultrafast universal RNA-seq aligner. Bioinformatics 29, 15–21 (2013).

15. Liao, Y., Smyth, G.K. & Shi, W. featureCounts: an efficient general purpose program for assigning sequence reads to genomic features. Bioinformatics 30, 923–930 (2014).

16. Love, M.I., Huber, W. & Anders, S. Moderated estimation of fold change and dispersion for RNA-seq data with DESeq2. Genome Biol 15, 550 (2014).

17. Fares, I. et al. Cord blood expansion. Pyrimidoindole derivatives are agonists of human hematopoietic stem cell self-renewal. Science 345, 1509–1512 (2014).

18. Fares, I. et al. EPCR expression marks UM171-expanded CD34. Blood 129, 3344–3351 (2017).

19. Ishitsuka, K., Nishikii, H., Kimura, T., Sugiyama-Finnis, A. & Yamazaki, S. Purging myeloma cell contaminants and simultaneous expansion of peripheral blood-mobilized stem cells. Exp Hematol 131, 104138 (2024).

20. Weinreb, C., Rodriguez-Fraticelli, A., Camargo, F.D. & Klein, A.M. Lineage tracing on transcriptional landscapes links state to fate during differentiation. Science 367 (2020).

21. Rodriguez-Fraticelli, A.E. et al. Single-cell lineage tracing unveils a role for TCF15 in haematopoiesis. Nature 583, 585–589 (2020).

22. Batista Napotnik, T., Polajzer, T. & Miklavcic, D. Cell death due to electroporation - A review. Bioelectrochemistry 141, 107871 (2021).

23. Dever, D.P. et al. CRISPR/Cas9 β-globin gene targeting in human haematopoietic stem cells. Nature 539, 384–389 (2016).

24. Huerga Encabo, H. et al. Loss of TET2 in human hematopoietic stem cells alters the development and function of neutrophils. Cell Stem Cell 30, 781–799 e789 (2023).

25. Nakauchi, Y. et al. The Cell Type-Specific 5hmC Landscape and Dynamics of Healthy Human Hematopoiesis and TET2-Mutant Preleukemia. Blood Cancer Discov 3, 346–367 (2022).

